# Filamentation of *Vibrio cholerae* is an adaptation for surface attachment and biofilm architecture

**DOI:** 10.1101/470815

**Authors:** Benjamin R. Wucher, Thomas M. Bartlett, Alexandre Persat, Carey D. Nadell

**Affiliations:** Department of Biological Sciences, Dartmouth College, Hanover, NH 03755, USA; Department of Microbiology and Immunology, Harvard Medical School, Boston, MA 02115; Institute of Bioengineering and Global Health Institute, School of Life Sciences, École Polytechnique Fédérale de Lausanne, Lausanne, Switzerland

## Abstract

Collective behavior in spatially structured groups, or biofilms, is the norm among microbes in their natural environments. Though microbial physiology and biofilm formation have been studied for decades, tracing the mechanistic and ecological links between individual cell properties and the emergent features of cell groups is still in its infancy. Here we use single-cell resolution confocal microscopy to explore biofilm properties of the human pathogen *Vibrio cholerae* in conditions closely mimicking its marine habitat. We find that some – but not all – pandemic isolates produce filamentous cells than can be over 50 μm long. This filamentous morphotype gains a profound competitive advantage in colonizing and spreading on particles of chitin, the material many marine *Vibrio* species depend on for growth outside of hosts. Furthermore, filamentous cells can produce biofilms that are independent of all currently known secreted components of the *V. cholerae* biofilm matrix; instead, filamentous biofilm architectural strength appears to derive from the entangled mesh of cells themselves. The advantage gained by filamentous cells in early chitin colonization and growth is counter-balanced in longer term competition experiments with matrix-secreting *V. cholerae* variants, whose densely packed biofilm structures displace competitors from surfaces. Overall our results reveal a novel mode of biofilm architecture that is dependent on filamentous cell morphology and advantageous in environments with rapid chitin particle turnover. This insight provides concrete links between *V. cholerae* cell morphology, biofilm formation, marine ecology, and – potentially – the strain composition of cholera epidemics.

## Introduction

Bacterial existence in the wild is predominated by life in spatially structured groups, termed biofilms (1, 2), which inhabit environments ranging from the rhizosphere (3), to chronic infections (4, 5), to the pipes of industrial and wastewater flow systems (6, 7), to the surfaces of marine snow (8–13). Although living in groups correlates with increased tolerance to many exogenous threats, including antibiotic exposure (14–16), biofilm-dwelling cells also experience intense competition for space and resources (2, 17, 18). Furthermore, cells in mature biofilms are generally non-motile and incur a trade-off between optimizing local competition versus dispersal to new environments (19–22). Balancing colonization, local growth, and dispersal is therefore a critical element of microbial fitness during biofilm formation, and understanding how bacteria have evolved to modulate this balance is a central challenge in microbial ecology. Here we study how variation in individual cell morphology impacts biofilm architecture and the competition/colonization tradeoff among different pandemic isolates of *Vibrio cholerae.*

*V. cholerae* is a notorious human pathogen responsible for the diarrheal disease cholera, but between epidemics, it persists as a common component of aquatic ecosystems, where it consumes chitin harvested from the exoskeletons of arthropods (23–25). To utilize this resource, *V. cholerae* must colonize and produce biofilms on chitin particles, creating a dense, resource-limited, and architecturally complex space for both inter-species and inter-strain competition (26–28). *V. cholerae* strains that are better adapted to colonize chitin surfaces, exploit the resources embedded in them, and spread to other particles are thus likely to be better represented in estuarine conditions. Although it remains poorly understood how V. *cholerae* makes the environmental-to-epidemic transition, emerging models implicate chitin-associated aggregates as disease vectors due to the high density of cells within them (24, 26, 29–33). Consequently, competition for access to space on chitin particles, and other substrates that form the basis for marine snow, may also impact which strains ultimately cause cholera pandemics (23, 34).

Many strains of *V. cholerae* can be found in the marine environment, and their relative abundances are continually in flux. This pathogen is characterized by its O-antigen, a glycan polymer on its outer membrane. While there are over two hundred O-antigen serotypes, only two have been known to cause major pandemics – serotype O1, comprised of both the Classical and El Tor biotypes, and serotype O139, which has arisen more recently alongside El Tor as the current leading cause of cholera in Southern Asia (35, 36). To better understand how competition to colonize biotic surfaces impacts the interactions between potentially co-occurring *V. cholerae* strains, we developed a microfluidic assay in which chitin particles are embedded in flow devices perfused with artificial sea water, and onto which cells could be readily inoculated to monitor colonization and biofilm growth.

We discovered that some pandemic isolates of *V. cholerae,* and in particular strain CVD112 of the O139 serogroup, filaments aggressively under nutrient-limited conditions, including on particles of chitin in sea water. Filamentation confers markedly altered chitin colonization and biofilm architecture relative to shorter cells. Differences in chitin colonization and biofilm architecture, in turn, strongly influence competition for space and resources, suggesting that normal-length and filamentous morphotypes are fundamentally adapted to different regimes of chitin particle turnover in the water column. Overall, our results highlight a novel mode of biofilm assembly and yield new insights into the fundamental roles of cell shape in the marine ecology of *V. cholerae.*

## Results

### Filamentation in conditions mimicking a marine habitat

We explored the biofilm architectures of different isolates of *V. cholerae,* focusing on N16961 (serogroup O1, El Tor biotype) and CVD112 (serogroup O139), representing the two major extant causative agents of the current cholera pandemic (36). The El Tor biotype has been used extensively in the study of biofilm formation and architecture (37–46); less is known about biofilms of the O139 serogroup (but see (47)). For visualization by microscopy, the far-red fluorescent protein-encoding locus *mKate2* (48) was inserted in single copy under the control of a strong constitutive promoter on the chromosome of CVD112 and its derivatives, while an analogous construct encoding the orange fluorescent protein *mKO-κ* (49) was inserted onto the chromosome of N16961 and its derivatives. We have shown previously that insertion of these fluorescent protein constructs to the chromosome does not substantially influence growth rate or other biofilm-associated phenotypes (27, 43).

In preliminary experiments, CVD112 and N16961 were inoculated into separate microfluidic devices and perfused with M9 minimal media containing 0.5% glucose. After 24 hours, confocal imaging revealed dramatic differences in the biofilm architecture of the two strains (Figure 1). Consistent with prior reports, N16961 grew in well-separated microcolonies of cells about 2 μm long, with each biofilm colony appearing to descend from one progenitor on the glass surface (Figure 1A) (42, 43). In contrast, CVD112 formed large groups of filamented cells entangled with one another and loosely associated with the glass surface (Figure 1B).

**Figure 1.**
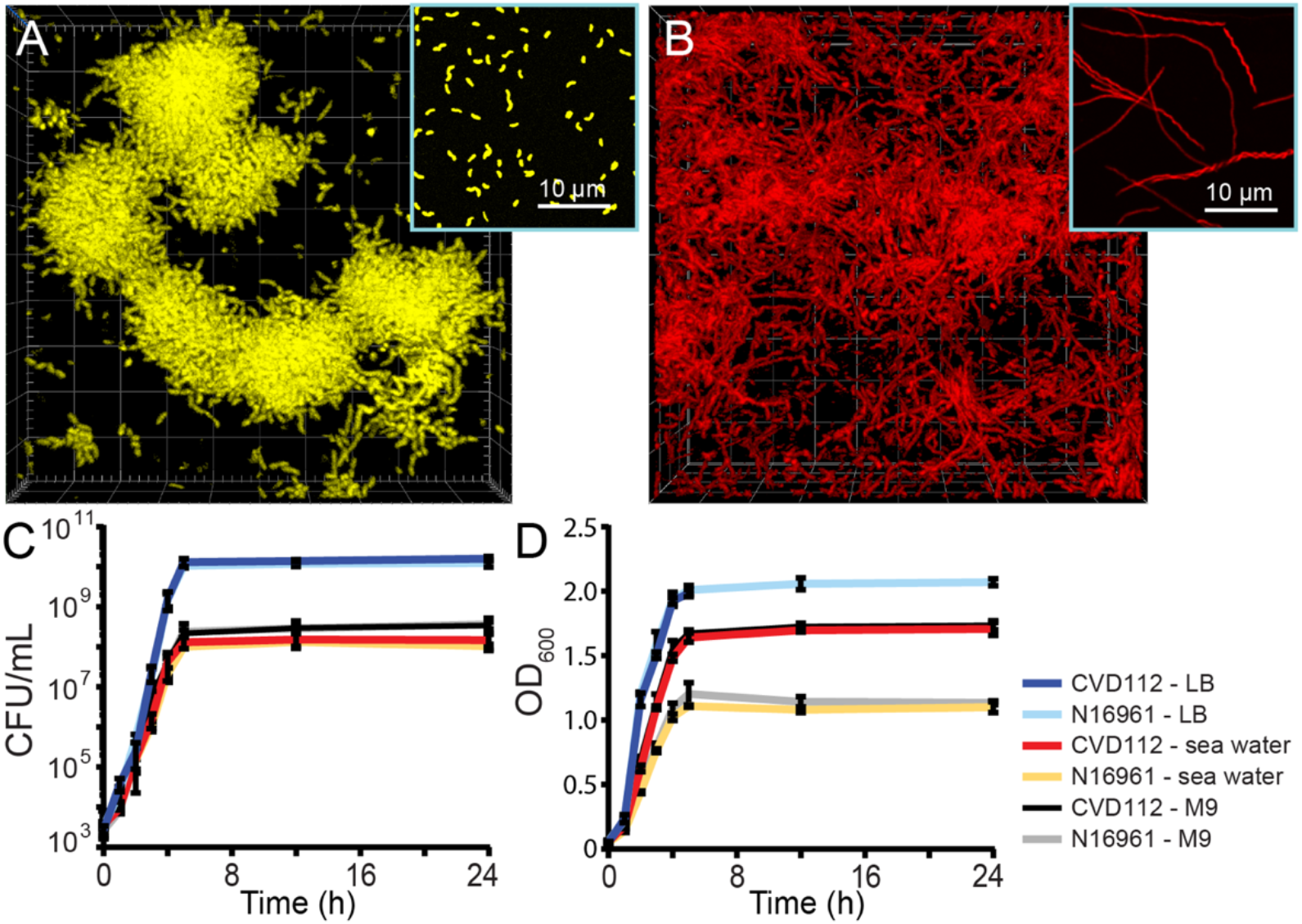
Cell morphology and biofilm structures of *Vibrio cholerae* N16961 (O1 serogroup) and CVD112 (O139 serogroup). (**A**) Cell cluster biofilms of El Tor str. N16961 (3-D render is 80×80×15μm [LxWxD], planktonic cells inset). (**B**) Filamented biofilms of O139 str. CVD112 (3-D render is 80×80×50μm [LxWxD], planktonic cells inset). Biofilm images in (**A**) and (**B**) were captured in glass-bottom chambers containing M9 minimal media with 0.5% glucose. Growth kinetics of CVD112 and N16961 in LB, artificial sea water with 0.5% GlcNAc, and M9 minimal media with 0.5% glucose, as measured by (**C**) colony-forming unit (CFU) count, and (D1) optical density at 600 nm (for each growth curve and condition, *n* = 3 biological replicates, each with 3 technical replicates).

We found that CVD112 filamentation occurs most strongly under nutrient-limited media environments, as CVD112 and N16961 cells were both observed exclusively as cells approximately 2 μm long when cultured in nutrient-rich lysogeny broth (LB) (Figure S1). In other circumstances, filamentation has been associated with an acute stress response prior to cell death, and since CVD112 filaments were seen in M9 media, but not LB, we first suspected that they might be in a disturbed physiological state. However, CVD112 showed no growth defect in either LB or M9 media relative to N16961 (Figure 1C).

In natural seawater environments, *V. cholerae* colonizes the exoskeletons of arthropods, where it consumes N-acetyl glucosamine (GlcNAc) released from chitin that *Vibrio* species digest by secretion of extracellular chitinases (26). We found that CVD112, but not N16961, filaments extensively in artificial seawater with 0.5% GlcNAc as the sole source of carbon and nitrogen (Figure S1). As was the case in LB and M9 media, CVD112 and N16961 growth curves are indistinguishable by CFU count in artificial seawater with GlcNAc, indicating similar rates of cell division (Figure 1C). Following this observation, we suspected that CVD112 produces more total biomass than N16961 per unit time in low-nutrient media (where CVD112 produces filaments, while N16961 produces cells of normal length), but not in high-nutrient media (where both strains produce cells of normal length). We tested this possibility by measuring the rate of total biomass production in liquid culture by optical density. As predicted, CVD112 produces biomass more quickly and to higher final density than N16961 in M9 media with glucose, and in artificial sea water with GlcNAc, but not in LB (Figure 1D). We visualized cells at regular time intervals over the course of their growth and confirmed that CVD112 begins filamenting in mid-exponential phase in sea water with GlcNAc, but not LB (Figure S1).

Filamentation of CVD112 in nutrient-limited conditions must derive in part from an increased cell elongation rate relative to cell division rate. This ratio is altered under nutrient-rich conditions such that CVD112 returns to a normal cell length, which we could observe by generating time-lapse image sets of CVD112 cells pre-filamented in low-nutrient media and placed under an agar pad made from LB. The filaments were found to septate rapidly, yielding progeny of the more conventionally observed 2 μm cell length for *V. cholerae* (Supplemental Movie 1). CVD112 thus appears to be healthy with respect to growth physiology, and it reversibly modulates its rates of elongation relative to division in low nutrient media so as to produce filaments. After inspecting other available isolates of *V. cholerae* in seawater with GlcNAc as the sole carbon and nitrogen source, we found that some strains of El Tor and O139 serogroups produce filaments in stationary phase, while others of each serogroup do not (Figure S2). Meanwhile, filamentation has previously been intimated by scanning electron microscopy in variants of MO10, another common model strain of *V. cholerae* O139 (50). Filamentation in seawater is thus found across a diversity of pandemic classes, and we hypothesized that filamentation is an adaptation – not specific to the O1 or O139 serogroups – that confers a fitness advantage in some natural conditions. As N16961 and CDV112 represent the strongest cases, respectively, of non-filamenting versus filamenting strains that we observed, we use these two for the remainder of the study as representative of their cell shape strategies in sea water conditions.

### Filamentation promotes colonization of chitin particles and enables matrix-independent biofilm formation

We next explored the biofilm morphologies of filamenting CVD112 and non-filamenting N16961 *V. cholerae,* simulating the natural conditions they experience in marine environments. We inoculated these strains in microfluidic channels decorated with traps to immobilize chitin particles (shrimp shell), a natural substrate of *V. cholerae* for biofilm growth and nutrient consumption (23, 28, 51). The chambers were perfused with artificial sea water lacking any additional source of carbon or nitrogen (27).

In the process of inoculating chitin with *V. cholerae* under flow, we noticed that CVD112 filaments were particularly adept at colonizing shrimp shell surface, with single cells often wound around the contours of individual chitin particles (Figure S3). This suggests that single filaments can undergo large deformation in shear flow, despite the stiffness of the cell wall, and that these flexible filaments can wrap around objects of similar or smaller length scale than the cell body. To determine if these properties give CVD112 a colonization advantage on chitin, we measured the attachment rates of filamentous CVD112 and non-filamenting N16961 by flowing a 1:1 mixture of the two strains (normalized by biomass per volume) onto chitin particles in artificial sea water. Filamentous *V. cholerae* does indeed colonize chitin more rapidly than short cells (Figure 2A). This qualitative result holds when the two strains’ colonization rates are tested in monoculture (Figure S4A), and when cells dispersing directly from previously occupied chitin particles are flowed into new chambers containing fresh chitin (Figure S4B).

**Figure 2.**
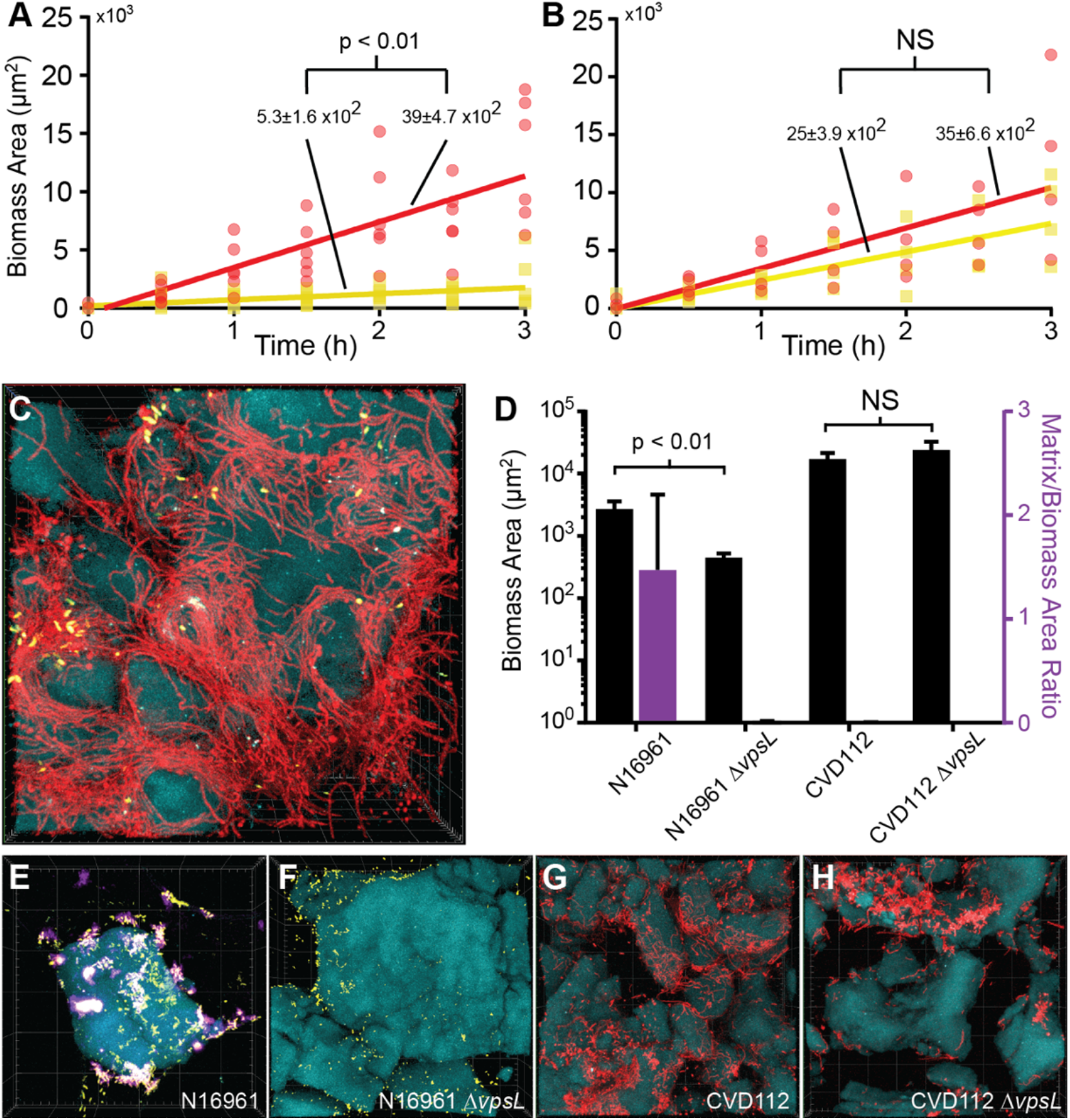
Filamentous *V. cholerae* CVD112 has an increased chitin colonization rate and produces matrix-independent biofilms on chitin in sea water. (**A**) The rates of CVD112 (red data) and N16961 (yellow data) accumulation onto fresh chitin particles in artificial sea water (*n* = 6 biological replicates). (**B**) As in (**A**), but here N16961 was pre-treated for 60 minutes with cefalexin, which causes it to filament in a manner similar to CVD112 without reducing cell viability (*n* = 4 biological replicates). (**C**) CVD112 cells (red) and N16961 (yellow) bound to pieces of chitin (blue), in artificial sea water (3-D render is 85×85×60μm [LxWxD]). (D) Mean biomass production (black bars, left vertical axis) and matrix normalized to biomass (purple bars, right vertical axis) for wild type and matrix-deficient *DvpsL* derivatives of CVD112 and N16961 (*n* = 3-6 biological replicates). (**E**) Wild Type N16961 (yellow), (**F**) N16961 *DvpsL* (yellow), (**G**) CVD112 (red) and (**H**) CVD112 *DvpsL* (red) on chitin (blue) in sea water. Matrix stain (Cy3-conjugated antibody to RbmA-FLAG) is shown in purple. 3-D renders in (**E**)-(**H**) are 175×175×40μm [LxWxD].

We next sought to determine whether the increased colonization ability of CVD112 was due to cell shape, as opposed to differences in other factors contributing to adhesion. To directly test the hypothesis that cell shape was responsible for increased chitin attachment, we treated N16961 cells with sub-inhibitory concentrations of cefalexin, which blocks cell division with minimal impact on cell viability or other aspects of *V. cholerae* morphology (52). N16961 pre-treated with cephalexin exhibited similar cell elongation and a chitin colonization rate statistically equivalent to that observed for CVD112 filaments (Figure 2B). These results indicate that filamentous morphology alone is sufficient to increase chitin surface colonization rate by an order of magnitude.

Following colonization, CVD112 produces biofilms that differ dramatically from those of N16961; they are composed of enmeshed cell filaments and lack the typical cell-cell packing and radial orientation associated with El Tor biofilm microcolonies (Figure 2C) (42, 44). The relative absence of tight cell-cell association prompted us to ask whether CVD112 was producing biofilm matrix, including *Vibrio* polysaccharide (VPS) and the adhesin RbmA, which interacts with the cell exterior and with VPS to hold neighboring cells in close proximity (40, 42–44, 53). To assess the contribution of matrix production to filamentous biofilms on chitin, we inoculated CVD112 and its isogenic *ΔvpsL* null mutant, which is unable to synthesize VPS or to accumulate any of the major matrix proteins (40). Biofilm production of wild type N16961, whose structure depends on matrix secretion (15, 40, 45), and its isogenic *ΔvpsL* null mutant were also measured for comparison. All strains contained a FLAG epitope inserted at the C-terminus of the native *rbmA* locus (40, 43). Fusion of a FLAG epitope to RbmA has been shown not to interfere with its function (40, 44), and allowed us to localize and quantify RbmA by immunostaining as a proxy for general matrix accumulation.

As expected, N16961 produced biofilms with abundant RbmA, while its isogenic *ΔvpsL* mutant was impaired for biofilm growth relative to the wild type parent and showed no matrix accumulation by RbmA staining (Figure 2D-F). The biomass and visible biofilm architecture of CVD112 and its *ΔvpsL* null mutant, on the other hand, were indistinguishable. CVD112 biofilms showed no detectable matrix accumulation, even in areas of dense growth (Figure 2D, G-H). Previous reports have suggested involvement of O-antigen and capsule polysaccharide in calcium-dependent biofilms of (non-filamentous) O139, but here we observed that filamentous biofilms did not rely on these factors: their biomass on chitin was unchanged in artificial sea water without calcium and in a *DwbfR* null background, which cannot synthesize O-antigen or capsule polysaccharide (Figure S5). The filamentous biofilm structures of CVD112 are thus independent of the currently known components of *V. cholerae* extracellular matrix. We infer that filamentous biofilm architecture can derive instead from entanglement of the cells themselves, which serve as both the actively growing biomass and the structural foundation of the community.

### *V. cholerae* filamentation is advantageous in frequently disturbed environments

Once we had discovered that filamentation provides *V. cholerae* with an advantage during chitin attachment, we sought to understand the relative fitness effects this cell morphology on chitin in co-culture with cells or normal length. To explore this question we inoculated CVD112 and N16961 together on chitin in artificial seawater, and their subsequent biofilm compositions were measured by confocal microscopy daily for 12 days.

When biofilms were left unperturbed for the full duration of the experiment, filamenting CVD112 cells had an initial advantage consistent with our colonization and growth rate experiments, but non-filamenting N16961 eventually increased in frequency to become the overwhelming majority of the population (Figure 3A,C). As noted above, filamentous CVD112 biofilms are independent of known matrix components, while N16961 secretes copious extracellular matrix within chitin-bound biofilms. Though CVD112 is superior in its initial surface occupation, its eventual displacement by N16961 is consistent with our prior work demonstrating that matrix secretion and cellcell packing confer a pronounced local competitive advantage within biofilms of *V. cholerae,* as well as resistance to spatial invasion by other bacteria (19, 20, 43, 54, 55). As opposed to matrix-replete biofilms of *V. cholerae,* biofilms of filamentous CVD112 permit competing strains to invade their interior volume: they do not maintain a grip over the space they initially claim during the colonization and early growth phases of the experiment (Figure S6).

**Figure 3.**
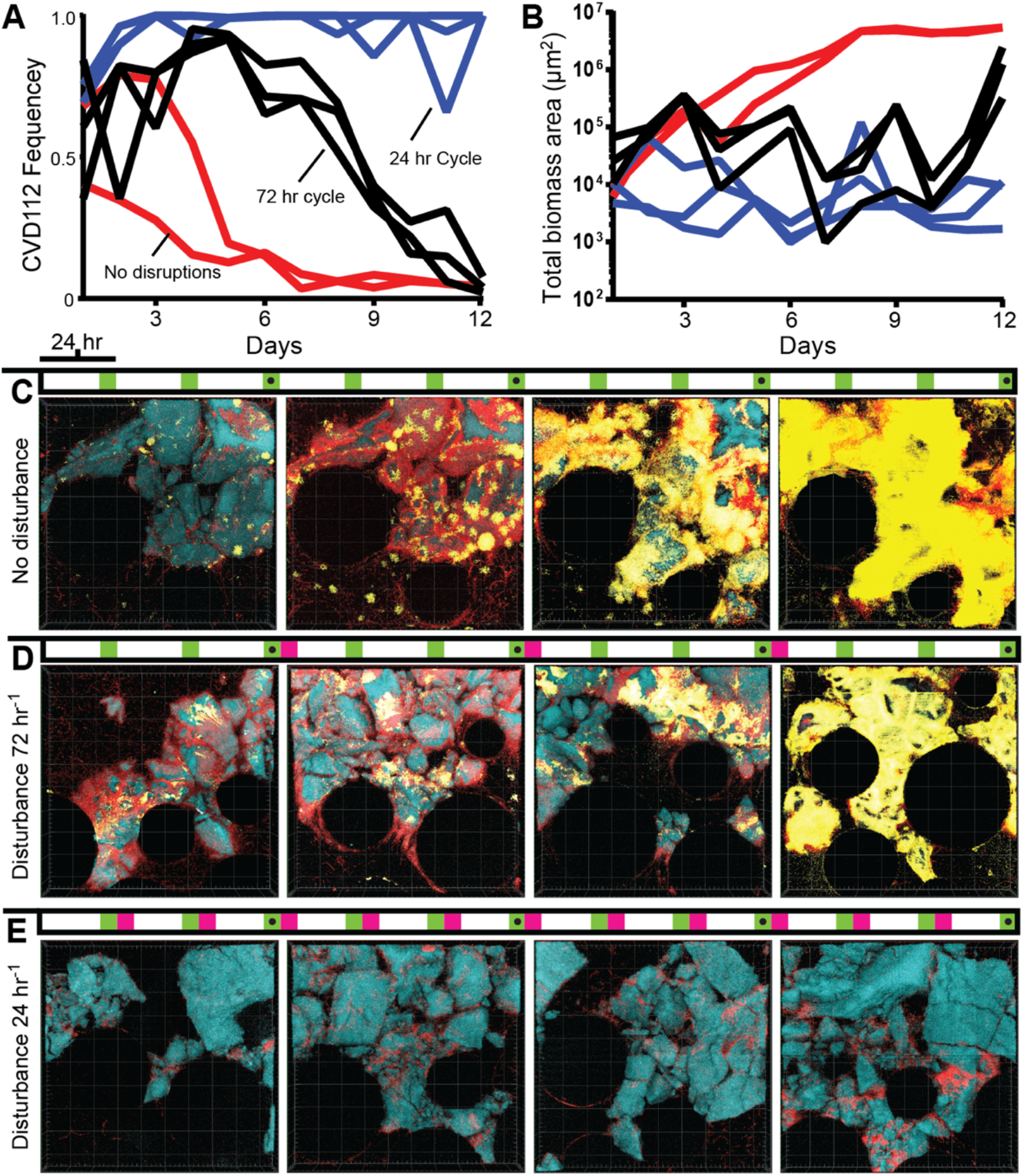
Competition between short cell N16961 (yellow) and filamenting CVD112 (red) on chitin (blue) in sea water. The two strains were grown together with different disturbance/recolonization regimes for 12 days. Chambers were either undisturbed (red traces in **A-B**, images in **C**), disrupted and reinoculated into new chitin chambers once per 72 hours (black traces in **A-B**, images in **D**), or disrupted and reinoculated into chambers every 24 hours (blue traces in **A-B**, images in **E**). Above each image series the treatment regime is shown with imaging times marked in green (representative images noted with black dots), and disturbance/recolonization events shown in magenta. All 3-D renders in panels C-E are 385×385×32μm [LxWxD].

Although matrix-producing N16961 outcompetes CVD112 within a patch of chitin-attached biofilm over time, the latter’s colonization advantage led us to speculate that it could be successful when residence times on a given chitin particle are shorter. This could be the case, for example, under frequently-agitated water column conditions or when chitin particle sizes are smaller, such that they are depleted quickly. To assess this idea, we repeated our competition experiments, but instead of tracking biofilm growth within a single microfluidic chamber over 12 days, we periodically used the liquid effluent exiting the chitin chamber to inoculate a new chamber containing fresh chitin, where a new competition would resume (Figure S7). This process simulates a disturbance event in which dispersal is advantageous and increases representation on new chitin particles elsewhere in the water column. We implemented two disturbance regimes, re-colonizing fresh chitin chambers once every 72 hours, or once every 24 hours.

When dispersed cells were taken from the effluent and allowed to recolonize fresh chitin every 72 hours, filamenting CVD112 cells showed a protracted early increase in frequency, but once again were eventually outcompeted by matrix-secreting, non-filamented N16961 cells (Figure 3A-B, D). However, when effluent collection and recolonization of fresh chitin occurred every 24 hours, the CVD112 strain dominated co-cultures for the full duration of the 12-day experiment (Figure 3A-B, E). The trajectories of these population dynamics were remarkably consistent from one run of the experiment to the next in all dispersal/recolonization regimes. This demonstrates the strength of the effects of competition during colonization, biofilm growth, and dispersal, relative to the influence of stochastic factors such as orientation of chitin particles in the chambers, or local variation in flow regime.

## Discussion

Individual cell morphology varies widely within and across bacterial species (56, 57), but in most cases it is not clear how cell shape relates to emergent structure and ecology of cell collectives such as biofilms. Here we have found that some isolates of *V. cholerae* produces long cell filaments under conditions closely matched to the natural marine environment. This cell morphology generates a pronounced advantage in chitin surface colonization and a matrix-independent biofilm architecture that permits rapid surface occupation, but also high dispersal rates. Filamentous cells’ superiority in early surface occupation, however, comes at a cost to long-term competition against other strains that invest in secretion of adhesive extracellular matrix. Filamentation is thus advantageous when patches of chitin turn over quickly, such that faster colonization and more easily reversible attachment are important components of fitness. Our results demonstrate that the shape of *V. cholerae* variants is a crucial factor controlling the relative investment into surface colonization, long-term biofilm robustness, and ease of dispersal back into the planktonic phase. These are fundamental elements of fitness for any biofilm-producing microbe, and they are especially important for the marine ecology of *V. cholerae* as it colonizes, consumes, and re-colonizes chitin in its environment outside of hosts.

Bacterial cell shape can serve a broad range of ecological functions (55–57). The curved shape of *Caulobacter crescentus,* for example, promotes the formation of biofilms as hydrodynamic forces reorient single cells to optimize daughter cell attachment (58); this process also nucleates clonal clusters under strong flow (59, 60). Simulations and experiments with engineered variants of *E. coli* suggest that rod-shaped bacteria can obtain a competitive advantage over spherical cells in colonies on agar plates, because rod-shaped cells burrow underneath spherical cells and spread more effectively to access fresh nutrients on the colony periphery (61). Filamentation has been observed in a wide variety of bacteria and eukaryotic microbes; this morphology is implicated in assisting spatial spread through soil or host tissue, and defense against phagocytosing ameboid predators (56, 62, 63). Here we have shown that filamentation can simultaneously alter surface colonization, biofilm architecture, and, as a result, the relative investment into rapid surface occupation versus long-term competitive success in a realistic environment.

The biophysical bases of our results are a topic of future work, but here we note that single filaments of *V. cholerae* can rapidly bend in shear flow, despite the stiffness of the bacterial cell wall (64). Our results demonstrate that sufficiently long bacteria can behave as elastic filaments (65); in analogy with the stretching behavior of polymers in flows that approach and split at the interface with an obstacle (i.e. extensional flows), we expect that filamentous bacteria experience shear that stretches them into alignment with stationary surfaces in the flow path (66–68). This process can increase the dwell time of filaments in proximity to obstacles in flow (Figure S8). We speculate that this process promotes attachment and wrapping of filaments around chitin particles (Figure S3), yielding a substantially augmented chitin colonization rate for filaments relative to shorter cells of *V. cholerae.*

Following surface attachment, filamentation allows the construction of biofilms in which cell-cell contacts generate a mesh network that is not dependent on currently known secreted components of the *V. cholerae* biofilm matrix. In this respect, filamentous biofilms may be analogous to a polymeric gel in which cell bodies are associated through physical entanglement rather than mutual attachment via secreted adhesives (69). Here, this cell network is more natively inclined to fast surface spreading and subsequent dispersal, but also porous and prone to physical invasion by competing strains or species. This strategy of rapid colonization and biomass accumulation but high reversibility of surface association is particularly well suited to fluctuating environments in which chitin particles are short-lived. This could be the case when particles are small and quickly consumed, or when disturbance events are common and destroy or disrupt chitin particles with high frequency.

The frequency of disturbance events and chitin particle size distribution are both likely to depend on the community ecology of planktonic organisms producing the chitin on which *V. cholerae* feeds in its aquatic environments. The preponderance and size of biofilm clusters has been strongly implicated in the ability of this pathogen to initiate infections that lead to epidemics: removing biofilm-like clusters from drinking water by filtration, for example, reduces the incidence of cholera infections by as much as 50% (70). Consequently, the differential competitive success of *V. cholerae* variants on chitin particles could be a significant determinant of which strains initiate disease outbreaks, which in turn often correlate with seasonal blooms of planktonic arthropod population growth (71). The size and shape of *V. cholerae* in its aquatic reservoir may therefore have consequences that emerge on large geographic scales in the extent and strain composition of cholera epidemics. More broadly, as we found that some but not all strains from both major pandemic serogroups produce filaments in seawater conditions, we expect that filamentation is a generic adaptation to rapid habitat turnover that may be found distributed quite widely in *Vibrio* spp. and other marine microbes for which particle surface attachment and biofilm growth are important fitness components.

## Materials and Methods

Methods for microfluidic device assembly, growth conditions, strain construction, growth curve measurements, septation imaging, biofilm cultivation, matrix staining, chitin colonization, competition assays, confocal microscopy, image analysis, and statistical analysis can be found further described in the SI appendix, Supplemental Materials and Methods.

## Author Contributions

CDN and BRW conceived the project. All authors contributed to experimental design. BRW performed strain construction, data collection, and image processing. BRW and CDN analyzed data and produced the figures. CDN and BRW wrote the manuscript with input from all authors.

## Acknowledgements

We are grateful to Rob McClung, Mary Lou Guerinot, Daniel Schultz, Karen Skorupski, Ryan Calsbeek, and Fitnat Yildiz for helpful comments on earlier versions of this manuscript, and to Kai Papenfort, Knut Drescher, Matthew Bond, and Swetha Kasetty for comments on the project. BRW is supported by a GANN Fellowship from Dartmouth College. AP is supported by the Swiss National Science Foundation (Projects grant 31003A_169377) and the Giorgio Cavaglieri Foundation. CDN is supported by the National Science Foundation (MCB 1817342), a Burke Award from Dartmouth College, a pilot award from the Cystic Fibrosis Foundation (STANTO15RO), and NIH grant P20-GM113132 to the Dartmouth BioMT COBRE.

## Supplementary Information

### Supplementary Materials and Methods

#### Strains and media

Supplementary Table S1 includes a full strain and plasmid list for this study. Strains are either derivatives of N16961 (El Tor serogroup) or CVD112 (O139 serogroup), excluding additional strains used for Figure S2. CVD112 and NT330 were generously given by the Skorupski lab in the Geisel School of Medicine at Dartmouth College. All modifications to N16961 and CVD112 were made using Escherichia coli S17-*λ*pir carrying the suicide vector pKAS32 for allelic exchange (72). All strains were grown in in LB, M9 minimal media with 0.5% glucose, or artificial sea water with either 0.5% GlcNAc or chitin as in (27). Antibiotics were used in the following concentrations: 100 μg/ml ampicillin, 50 μg/ml kanamycin, 50 μg/ml polymyxin B, 1000 μg/ml streptomycin and 5 μg/ml cefalexin. All chemicals and reagents were purchased from Millipore Sigma unless otherwise stated.

#### Plasmid Construction

All restriction enzymes and ligase were purchased from New England biolabs, and PCR reagents were purchased from BioRad. Codon optimized versions of *mKO-_κ_* and *mKate2* were purchased from Invitrogen. pCN764 (for insertion of *mKO-_κ_* to the *lacZ* locus) and pCN765 (for insertion of *mKate2* to the *lacZ* locus) were constructed by amplification of the flanking regions upstream and downstream of the lacZ open reading frame and fusing the respective genes for these fluorescent proteins to a synthetic Ptac for high expression from a single chromosomal locus. The lacZ-flanking fragments and fluorescent protein expression constructs were then combined using overlap extension PCR. These conjoined fragments were cloned into the pKAS32 vector backbone of pCN251 using the enzymes BsrG1 and BspE1 and ligated with T4 DNA ligase. These constructs were then introduced into Escherichia coli S17-*λ*pir by electroporation and conjugated into *V. cholerae.* Deletions of *vpsL, rbmA* and *wbfR* were made by cloning 1kb sequences up- and down-stream of the respective reading frames, joining these by PCR overlap extension, and cloning these fused products into the pKAS32 backbone. Insertion of the 3xFLAG epitope to the C-terminus of *rbmA* was performed using pKAS32-based allelic exchange as described previously (43).

#### Liquid growth curve experiments

*V. cholerae* strains were grown at 37° C shaking in LB overnight prior to the experiment. Overnight cultures were then back diluted to an OD_600_ of 0.01 in LB, M9 minimal medium with 0.5% glucose, or artificial sea water with 0.5% GlcNAc. OD_600_ were monitored every hour and once overnight. To measure viable cell count, serial dilutions were also performed at each timepoint and plated on LB agar for colony forming units. Each growth curve was performed with three biological replicates from independent overnight cultures and three technical replicates per biological replicate. Measurements for optical density were taken with a CO8000 cell density meter (Biochrom, Cambridge UK).

#### CVD 112 septation imaging

CVD 112 was grown overnight in sea water with 0.5% GlcNAc. Thirty minutes prior to imaging, 1 ml of overnight culture was spun down, resuspended in fresh LB, and incubated at 37° C to encourage septation. 10 μl of this culture was then spotted onto a #1.5 cover glass and covered with an LB agar pad. Phase contrast images were taken every 5 minutes at 100x magnification and a video was rendered using Nikon NIS elements (Tokyo, Japan).

#### Microfluidic device assembly

The microfluidic devices used consist of poly-dimethylsiloxane (PDMS) bonded to size # 1.5 36mm X 60mm cover glass (ThermoFisher, Waltham MA) using standard soft lithography techniques (73, 74). Two designs of chamber were used, planar and columnar. To establish flow in these chambers, media was loaded into 1ml BD plastic syringes with 25-gauge needles. These syringes were joined to #30 Cole palmer PTFE tubing (inner diameter 0.3mm), which was connected to pre-bored holes in the microfluidic device. Tubing was also placed on the opposite end of the chamber to direct the effluent to a waste container. Syringes were mounted to syringe pumps (Pico Plus Elite, Harvard Apparatus), and flow was maintained at 0.2ul min^-1^ for all experiments.

#### Biofilm growth and matrix staining on chitin

Prior to bacterial inoculation, chitin flakes were be sterilized with 70% ethanol and washed in sea water (as in Drescher et al. (27)). These flakes were suspended in sea water and flowed into a columnar chamber at high speed using a 1 ml syringe attached to a small length of PTFE tubing. After 30 minutes, sea water was introduced into the device at a rate of 0.2ul/min. Overnight cultures of each strain were normalized to an OD_600_ of 1.0, inoculated into a microfluidic chamber previously filled with chitin, and allowed to rest for 30 minutes. The devices were then run at room temperature for different periods of time depending on the experimental design. For experiments in which RbmA matrix protein was stained for localization and quantification: twelve hours prior to imaging, the influent media was replaced with sea water supplemented with 1ug/ml anti-FLAG antibody conjugated to the fluorescent dye Cy3 (Millipore-Sigma) to stain FLAG-tagged RbmA protein *in situ.*

#### Chitin colonization experiments

Strains were grown overnight at 37°C in sea water so that CVD112 would be filamented prior to the experiment. In Figure 2B, strain N16961 was sub-cultured, grown to mid log phase and exposed to a sub inhibitory concentration of cefalexin (5ug/ml) to induce filamentation as in Bartlett et al. (52). Cultures were then diluted to an OD_600_ of 0.1. equal amounts of each strain were mixed 1:1 in a BD 1ml plastic syringe (for experiments in Figure S4, strains were loaded at an OD_600_ of 0.1 to syringes in monoculture). The syringes were connected to a new chamber filled with chitin flakes. These chambers were then perfused with the corresponding culture preparations at a flow rate of 0.2μl/min and imaged every 30 minutes for 3 hours.

#### Chitin competition assays

Microfluidic devices were prepared with chitin as described above. Fluorescent protein expressing strains of CVD112 and N16961 (producing mKate2 and mKO-κ, respectively) were grown in LB overnight at 37°C. These cultures were then equalized to an OD_600_ of 1, mixed 1:1 and inoculated into the microfluidic device. After 30 minutes, flow of fresh sea water lacking any carbon or nitrogen source were introduced at 0.2 μl/min. Under the no disruption condition, the devices were run at room temperature for 12 days and imaged daily. In conditions with disruption events, new microfluidic devices containing chitin were prepared, and the effluent from the currently incubating device was flowed into the new devices for 30 min. After colonization of the new device, the previous device was discarded. Between these disruption events, chambers were incubated at room temperature and imaged daily (see Figure S6).

#### Microscopy and image analysis

Biofilms inside microfluidic chambers were imaged using a Zeiss LSM 880 confocal microscope with a 40x / 1.2 N.A. water objective. A 543-nm laser line was used to excite mKO-_ĸ_, and a 594-nm laser line was used to excite mKate2. A 458-nm laser line was used to excite chitin autofluorescence. For biomass accumulation and competition experiments, images were acquired as a wide view tile scan, so that all chitin in each chamber would be present for analysis. In experiments involving RbmA-FLAG staining, representative images were taken at several locations within the chamber and averaged per biological replicate. Figure panels were rendered using Zeiss ZEN blue software. To quantify bacterial or matrix biomass per area, we used customized scripts in MATLAB (MathWorks, Natick, MA) as in Drescher et al. 2014 (27) and Nadell et al. (43).

#### Statistics

All statistical analyses were performed in GraphPad prism. All reported comparisons are Wilcoxon signed ranks tests with Bonferroni correction. For biomass accumulation experiments reported in Figure 2A-B, individual slopes were determined via a linear regression for each biological replicate. These slopes were used as data points for the corresponding comparison tests.

### Supplementary Figures

**Figure S1.**
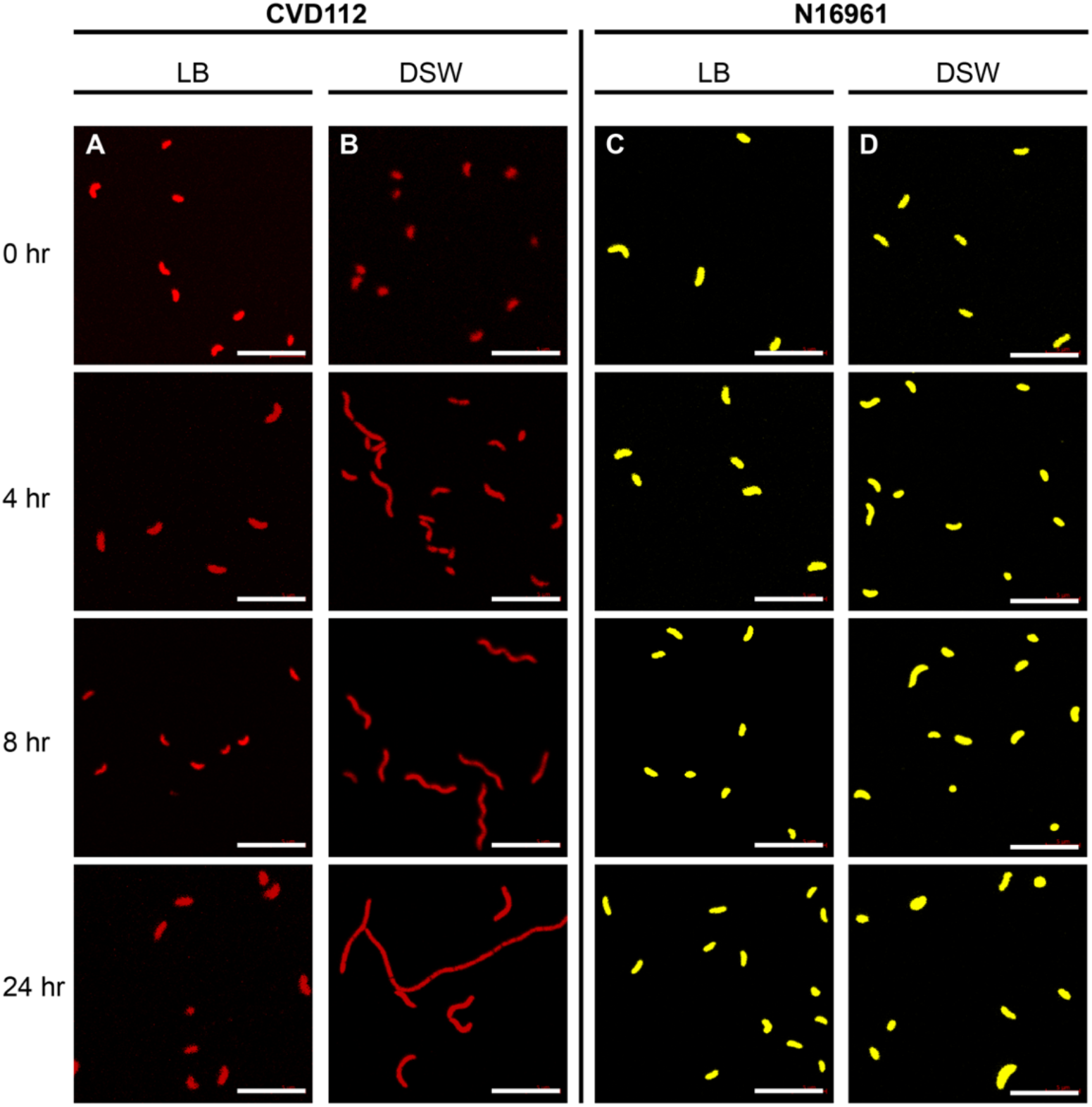
Snapshots of aliquots of CVD112 cells (red, left) and N16961 cells (yellow, right) over the course of planktonic growth curve experiments in LB or artificial sea water with 0.5% GlcNAc. CVD112 filaments in sea water, but not LB, while N16961 does not filament in either condition. Scale bars denote 10 μm.

**Figure S2.**
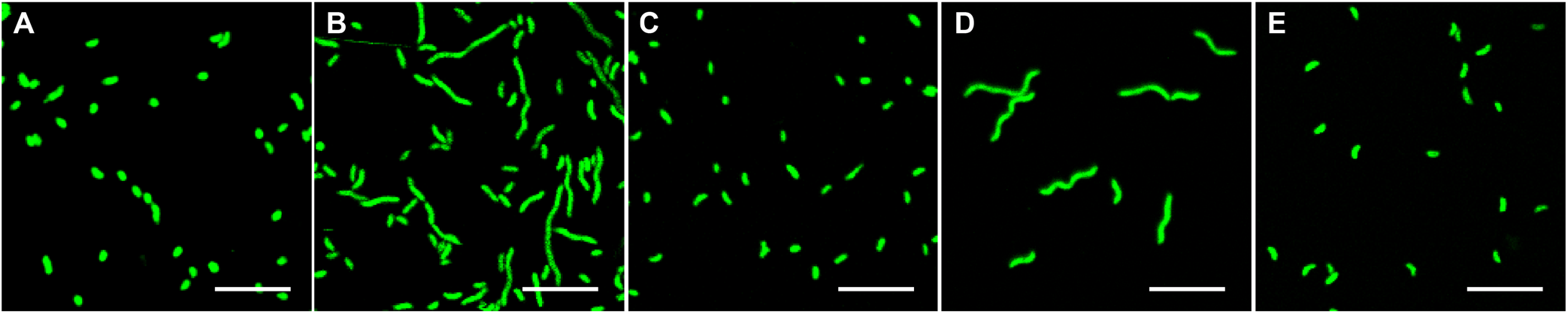
Fluorescence images of *V. cholerae* strains. (**A**) O1 - El Tor, N16961, (**B**) O1 - El Tor, C6706, (**C**) O1 – Classical, RT4633 (**D**) O139 - CVD112, and (E) O139 - NT330, taken in stationary phase after growth in artificial sea water with 0.5% GlcNAc as the sole source of carbon and nitrogen. Cells were stained with Syto 9. Scale bars denote 10 μm.

**Figure S3.**
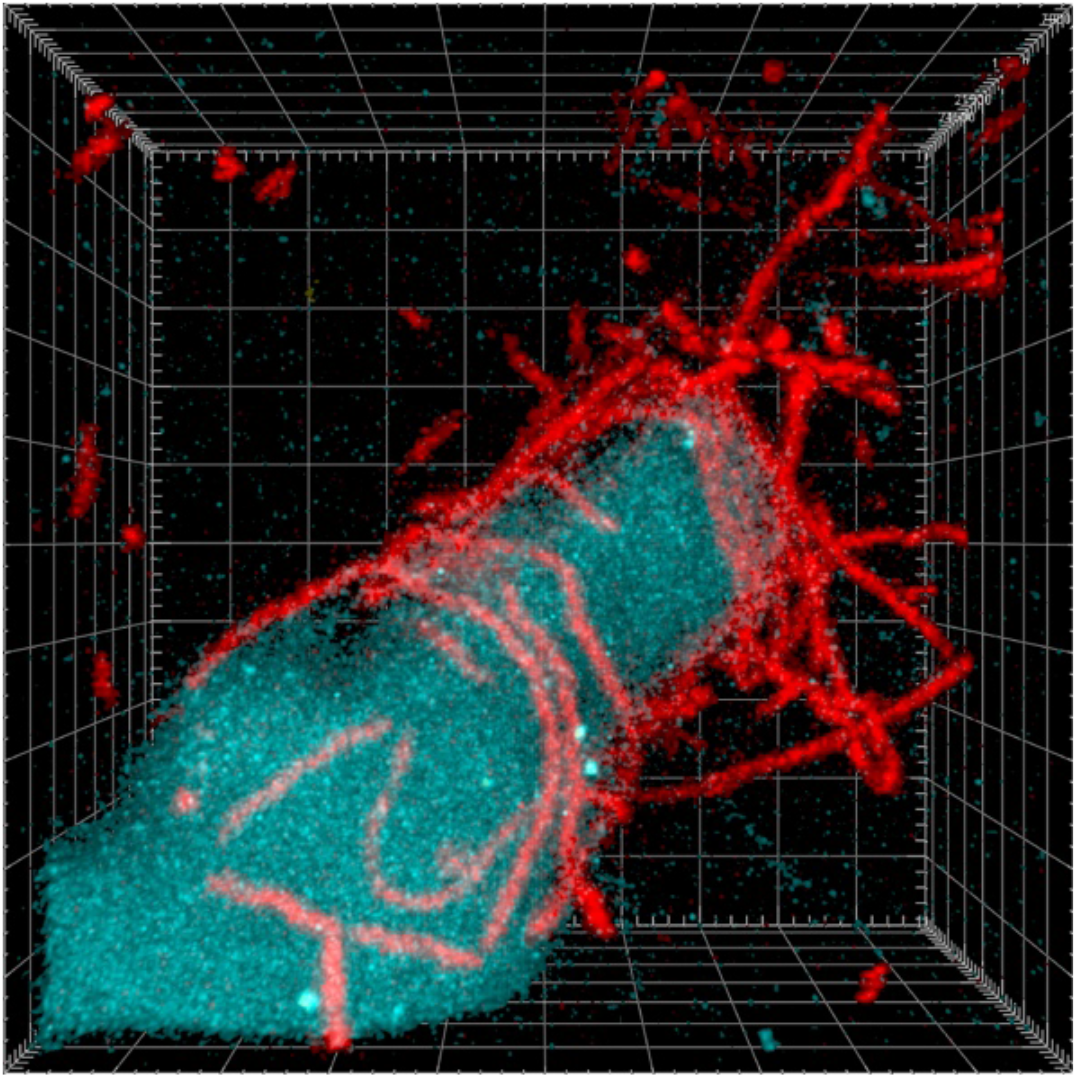
Filamentous cells of *V. cholerae* CVD112 (red) wrapped around the contours of a chitin particle (blue) in artificial sea water. This 3-D rendering is 35×35×60μm [LxWxD].

**Figure S4.**
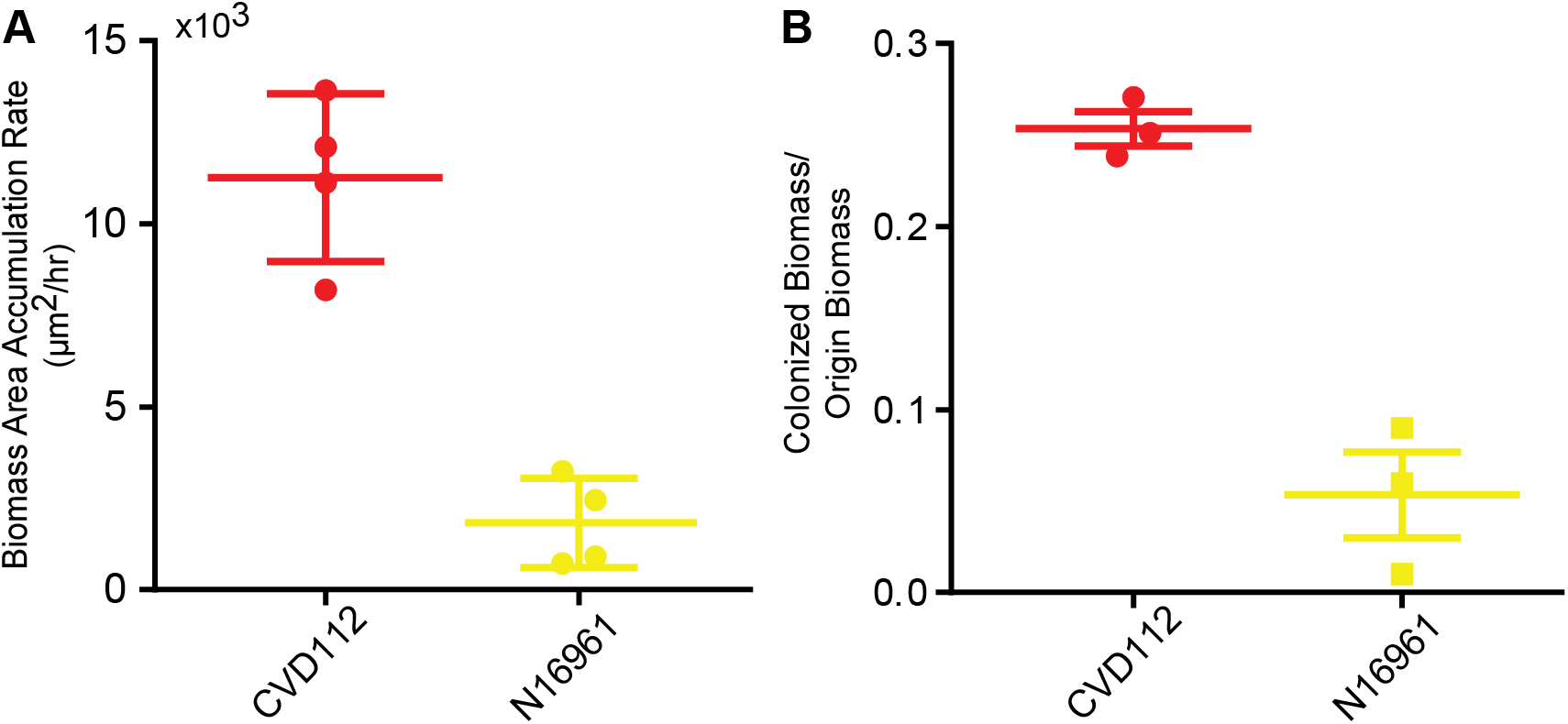
Chitin colonization rates of filamentous *V. cholerae* CVD112 and non-filamentous N16961. (**A**) CVD112 and N16961 were flowed in monoculture into chambers containing fresh chitin particles, and their biomass accumulation rates (biomass area per hr) were measured (*n* = 4 biological replicates). (**B**) Chambers containing monoculture biofilms of CVD112 or N16961 growing on chitin particles were allowed to grow for 48 hr, after which the effluent from these chambers was connected to the fluid inlet of chambers containing fresh chitin for 1 hr (*n* = 3 biological replicates). The total biomass area of cells colonized on the fresh chitin particles was measured and normalized to the biomass of cells present in the previous chamber from which effluent was collected for the colonization experiment.

**Figure S5.**
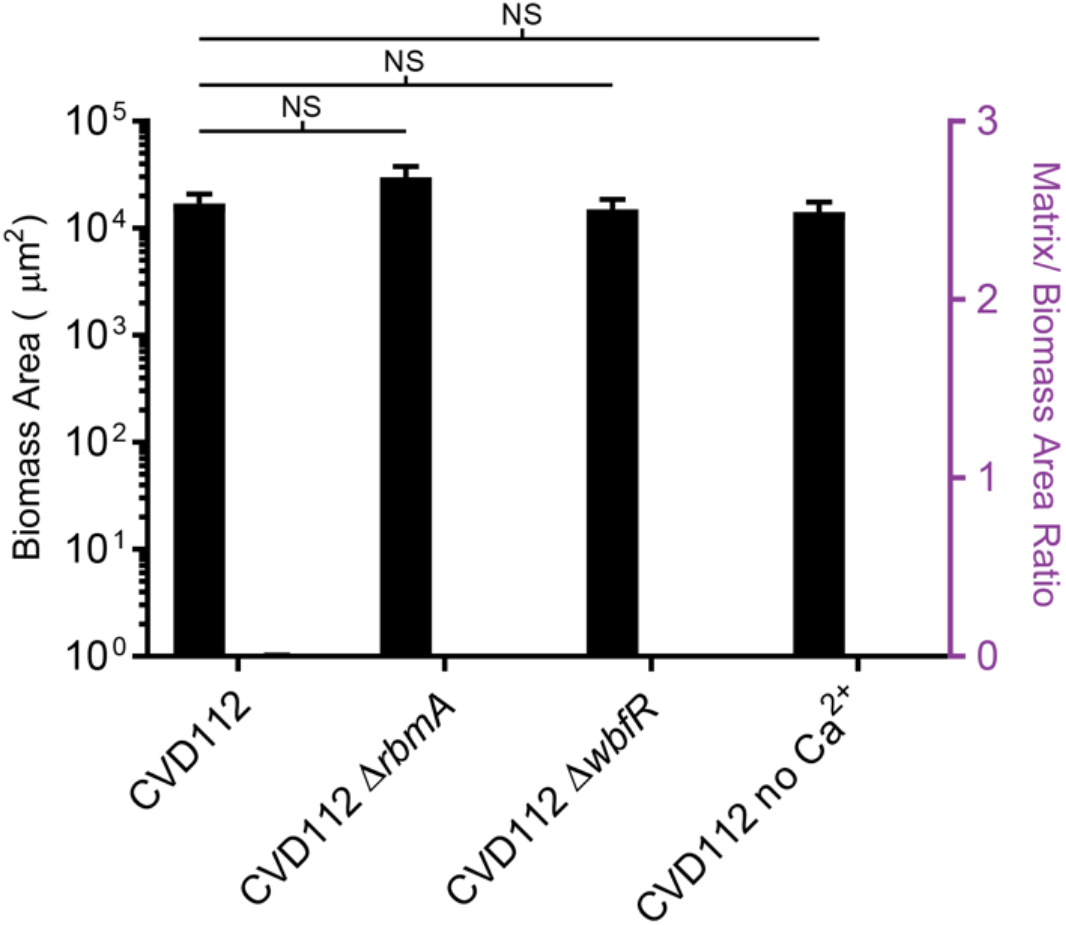
Filamentous biofilms of CVD112 were grown on chitin, with the parental wild type shown at left for comparison. No difference in biomass accumulation was observed for biofilms lacking the cell-cell adhesin RbmA (*n* = 4 biological replicates). Biofilms of O139 have been reported to rely on calcium and a combination of O-antigen and capsular polysaccharide, but here we found that *wbfR* null mutants (which cannot synthesize O-antigen or capsule) are not impaired for filamentous biofilm growth, nor are wild type CVD112 biofilms grown in sea water without Ca^2+^ (*n* = 4 biological replicates).

**Figure S6.**
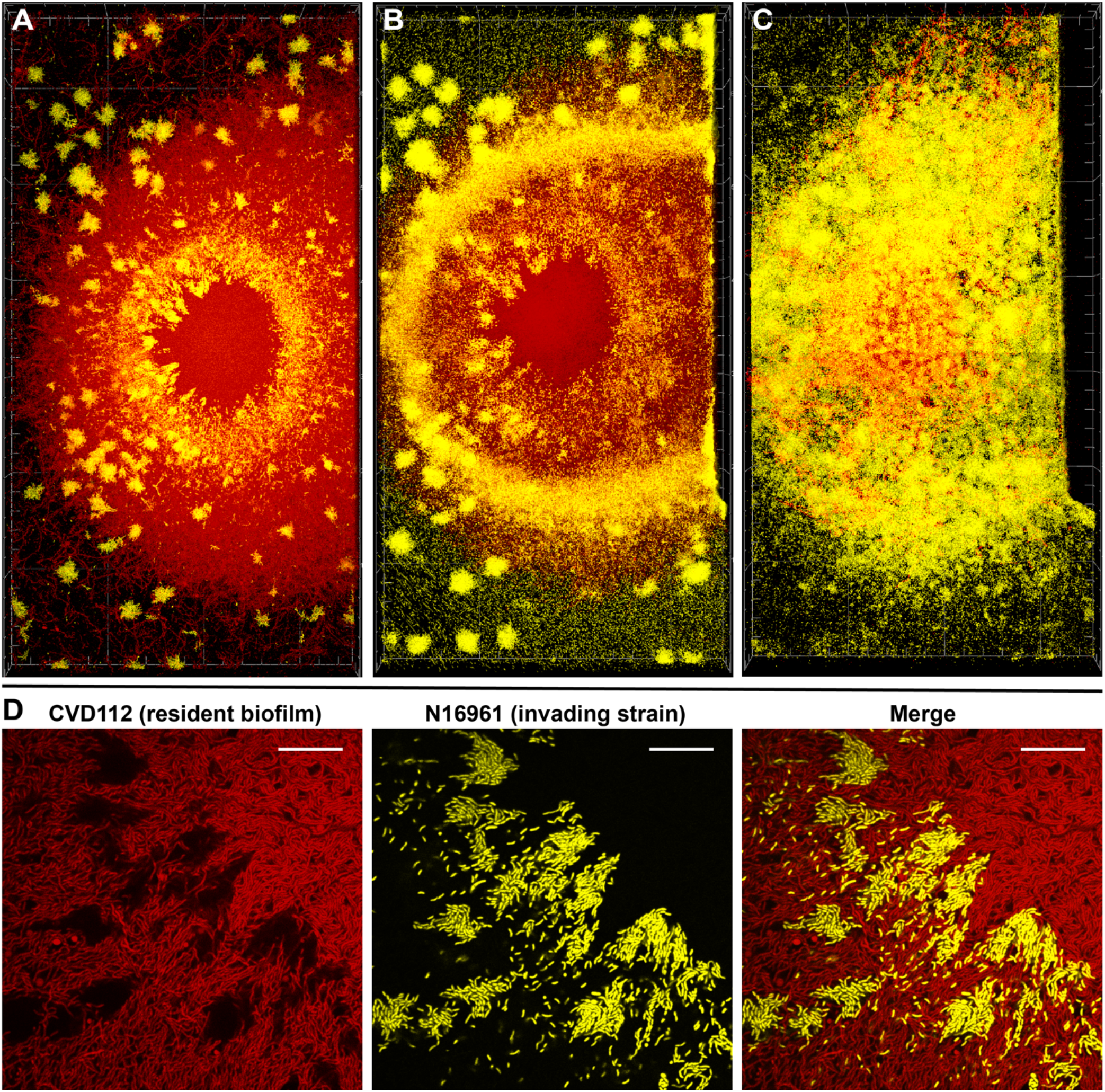
Filamentous biofilms of CVD112 (red) can be invaded and physically displaced by cells of N16961 (yellow). Here biofilms of CVD112 were grown in isolation, and subsequently exposed to cells of N1691. The progression of the invasion is shown at (**A**) 24, (**B**) 48, and (**C**) 72 hours after invading N16961 cells were introduced to the system. The 3-D renders in A-C are 675×350×45 μm [LxWxD]. (**D**) A magnified portion of the biofilms from panel B, showing that the invading N16961 cells displace CVD112 from the substratum on which they are growing. Scale bars denote 20 μm.

**Figure S7.**
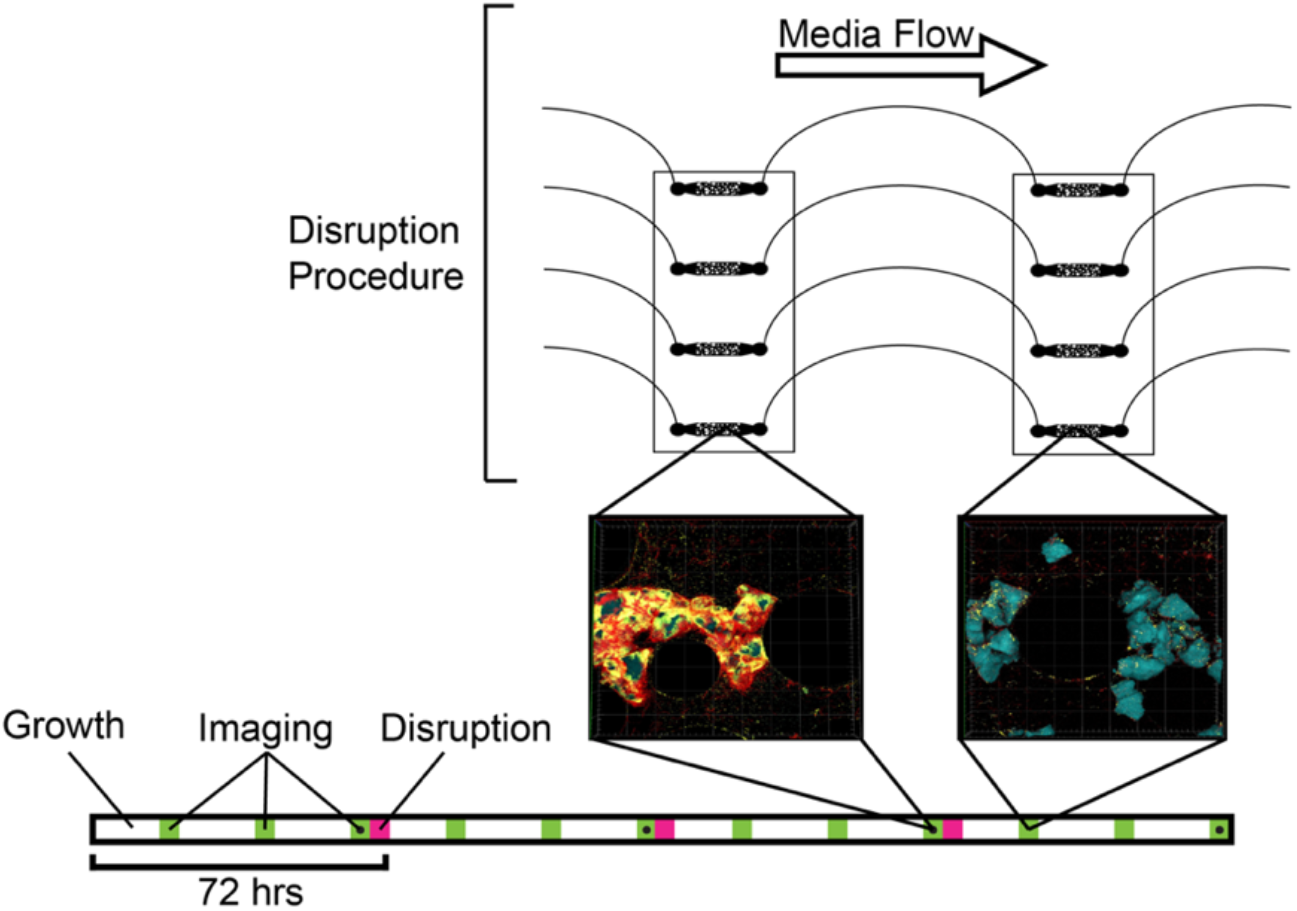
An illustration of the regime for competition experiments with disturbance/recolonization. *V. cholerae* CVD112 and N16961 were introduced in co-culture to microfluidic devices containing chitin and imaged daily (imaging points are indicated by green square events in the timeline above). Every 72 hours (magenta square events), the effluent from the current chamber was used to inoculate a new chamber containing fresh chitin, which was then monitored daily until the next disturbance event. In another treatment (see Figure 3, main text), disturbance/recolonization events were performed every 24 hours.

**Figure S8.**
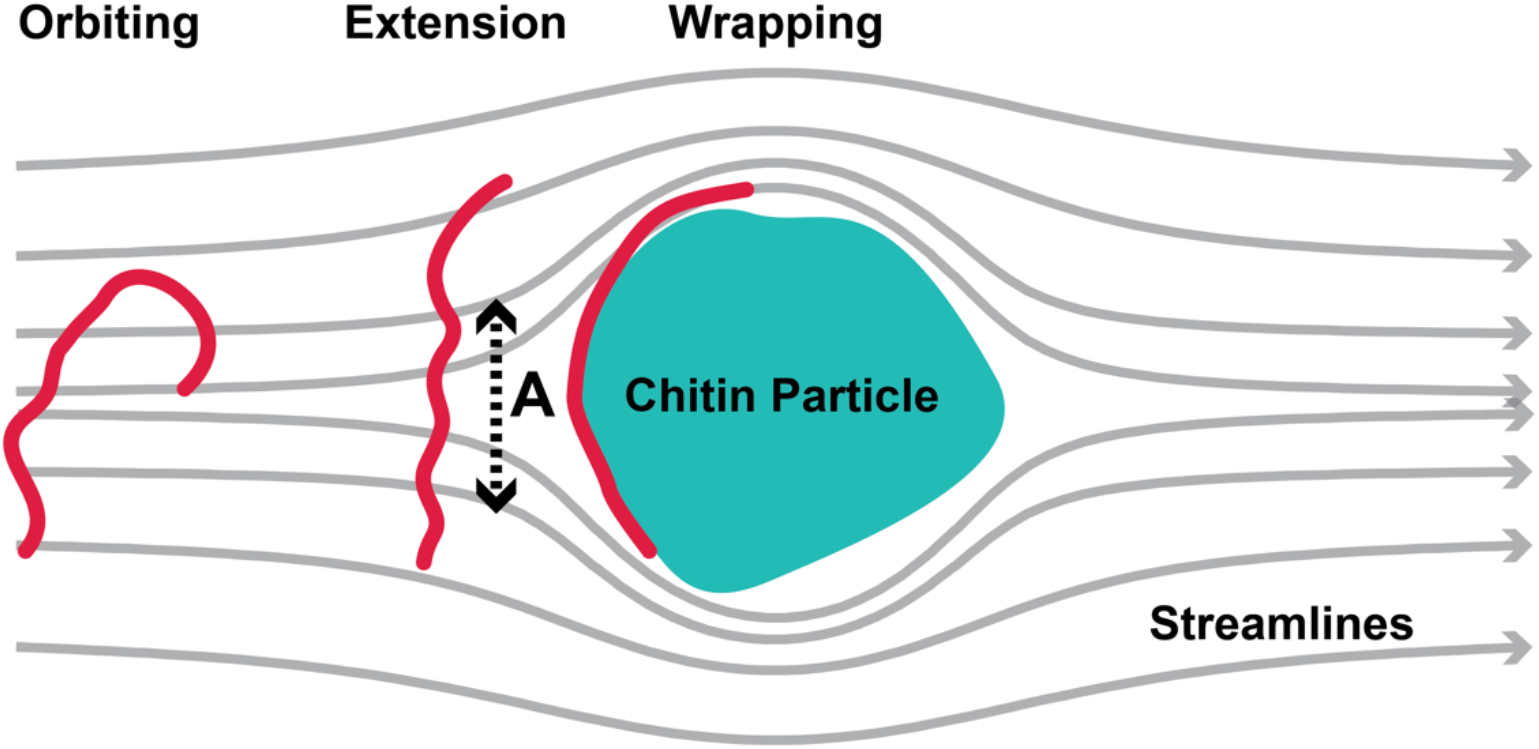
A fluid dynamics model for cell filament stretching in flow near chitin particles. The flow profile near the surface of a cylinder can be described as an extensional flow. This description is most accurate close the particle surface at the stagnation point (where streamlines split, point A above). Elastic filaments such as polymers experiencing this extensional flow align in the direction of extension. We therefore anticipate that single filamentous *V. cholerae* cells align with approaching chitin particles in flow. In addition, extended cells span either side of the stagnation point, effectively increasing their residence time near the particle, while short cells ballistically follow streamlines. Altogether, stretching and increased residence time may participate to the improved ability of filamentous cells to encounter and attach to chitin particles.

**Supplementary Table 1:** *V. cholerae* strains and plasmids.

| Strain | Relevant markers/ genotypes | Source |
| --- | --- | --- |
| *E. coli* |  |  |
| S17-1λpir | wild type | (75) |
| *V. cholerae* |  |  |
| CNV49 | wild type strain N16961 (O1 El tor Sm resistant) | (26) |
| CNV116 | N16961 *rbmA*-3xFLAG, *lacZ*:P<sub>tac</sub>-*mKate2* | This study |
| CNV121 | N16961 *rbmA*-3xFLAG, *lacZ*:P<sub>tac</sub>-*mKO-κ* | This study |
| CNV203 | N16961 Δ*vpsL*, *rbmA*-3xFLAG, *lacZ*:P<sub>tac</sub>-*mKate2* | This study |
| CNV180 | CVD112 (RT4102 O139 Sm resistant at 1mg/ml) | (76) |
| CNV184 | CVD112 *lacZ*:P<sub>tac</sub>-*mKate2* | This study |
| CNV 202 | CVD112 *rbmA*-3xFLAG, *lacZ*:P<sub>tac</sub>-*mKate2* | This study |
| CNV205 | CVD112 Δ*vpsL*, *rbmA*-3xFLAG, *lacZ*:P<sub>tac</sub>-*mKate2* | This study |
| CNV206 | CVD112 Δ*wbfR*, *rbmA*-3xFLAG, *lacZ*:P<sub>tac</sub>-*mKate2* | This study |
| CNV 207 | CVD112 Δ*rbmA* *lacZ*:P<sub>tac</sub>-*mKate2* | This study |

| Plasmid | origin, marker | comments | templates, primers | source |
| --- | --- | --- | --- | --- |
| pCN251 | pR6K, Amp, Kan | pKAS32 |  | (72) |
| pCN764 | pR6K, Amp, Kan | pKAS32 for insertion of *lacZ*:P<sub>tac</sub>-*mKO-κ* | Cno233-236 | This study |
| pCN765 | pR6K, Amp, Kan | pKAS32 for insertion of *lacZ*:P<sub>tac</sub>-*mKate2* | cno233-236 | This study |
| pCN767 | pR6K, Amp, Kan | pKAS32 Δ*wbfR* | cno302-305 | This study |
| pCN006 | pR6K, Amp, Kan | pKAS32 Δ*vpsL* | | C. M. Waters, Michigan State University, East Lansing, MI |
| pCN098 | pR6K, Amp | pKAS32 Δ*rbmA* | | (43) |
| pCN189 | pR6K, Amp | pKAS32 *rbmA*-3xFLAG insertion |  | (43) |

**Supplementary Table 2:**
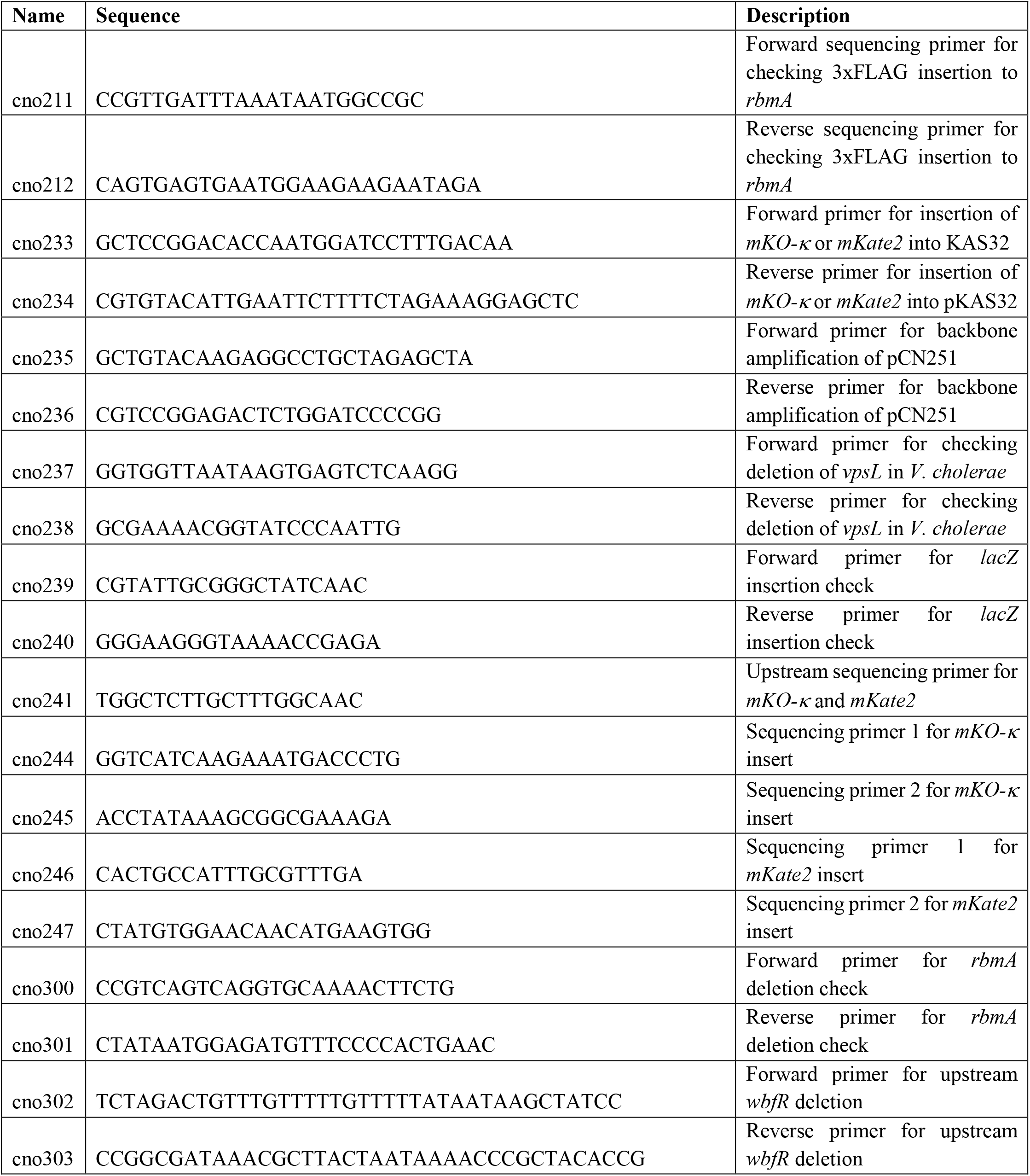
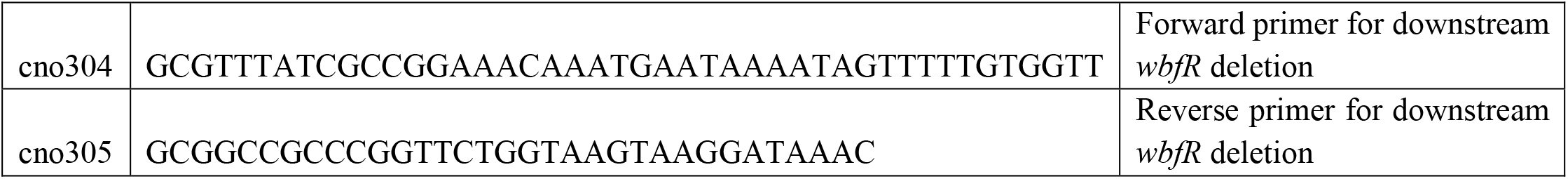
Primers used in plasmid construction.

## References

1. Flemming H-C, et al. (2016) Biofilms: an emergent form of bacterial life. Nat Rev Microbiol 14(9):563–575.

2. Nadell CD, Drescher K, Foster KR (2016) Spatial structure, cooperation, and competition in bacterial biofilms. Nat Rev Microbiol 14:589–600.

3. Rudrappa T, Biedrzycki ML, Bais HP (2008) Causes and consequences of plant-associated biofilms. FEMS Microbiol Ecol 64(2):153–166.

4. Ciofu O, Hansen CR, Hoiby N (2013) Respiratory bacterial infections in cystic fibrosis. Curr Opin Pulm Med 19(3):251–258.

5. Wolcott R (2015) Disrupting the biofilm matrix improves wound healing outcomes. J Wound Care 24(8):366–371.

6. Barnes RJ, et al. (2015) Nitric oxide treatment for the control of reverse osmosis membrane biofouling. Appl Environ Microbiol 81(7):2515–2524.

7. Yuki Miura, Yoshimasa Watanabe and, Okabe* S (2006) Membrane Biofouling in Pilot-Scale Membrane Bioreactors (MBRs) Treating Municipal Wastewater: Impact of Biofilm Formation. doi:10.1021/ES0615371.

8. Cottingham KL, Chiavelli DA, Taylor RK (2003) Environmental microbe and human pathogen: the ecology and microbiology of Vibrio cholerae. Front Ecol Environ 1(2):80–86.

9. Dang H, Lovell CR (2016) Microbial Surface Colonization and Biofilm Development in Marine Environments. Microbiol Mol Biol Rev 80(1):91–138.

10. Cordero OX, Ventouras LA, DeLong EF, Polz MF (2012) Public good dynamics drive evolution of iron acquisition strategies in natural bacterioplankton populations. Proc Natl Acad Sci U S A 109(49):20059–20064.

11. Datta MS, Sliwerska E, Gore J, Polz MF, Cordero OX (2016) Microbial interactions lead to rapid micro-scale successions on model marine particles. Nat Commun 7:11965.

12. Enke TN, Leventhal GE, Metzger M, Saavedra JT, Cordero OX (2018) Microscale ecology regulates particulate organic matter turnover in model marine microbial communities. Nat Commun 9(1):2743.

13. Leventhal GE, et al. (2018) Strain-level diversity drives alternative community types in millimetre-scale granular biofilms. Nat Microbiol 3(11):1295–1303.

14. Hoiby N, Bjarnsholt T, Givskov M, Molin S, Ciofu O (2010) Antibiotic resistance of bacterial biofilms. Int J Antimicrob Agents 35(4):322–332.

15. Vidakovic L, Singh PK, Hartmann R, Nadell CD, Drescher K (2018) Dynamic biofilm architecture confers individual and collective mechanisms of viral protection. Nat Microbiol 3:26–31.

16. Mah T-FC, O’Toole GA (2001) Mechanisms of biofilm resistance to antimicrobial agents. Trends Microbiol 9(1):34–39.

17. Hibbing ME, Fuqua C, Parsek MR, Peterson SB (2010) Bacterial competition: surviving and thriving in the microbial jungle. Nat Rev Micro 8(1):15–25.

18. Nadell CD, Xavier JB, Foster KR (2009) The sociobiology of biofilms. Fems Microbiol Rev 33(1):206–224.

19. Nadell CD, Bassler BL (2011) A fitness trade-off between local competition and dispersal in Vibrio cholerae biofilms. Proc Natl Acad Sci USA 108(34):14181–14185.

20. Yan J, Nadell CD, Bassler BL (2017) Environmental fluctuation governs selection for plasticity in biofilm production. ISME J 11(7):1569–1577.

21. Yawata Y, et al. (2014) Competition-dispersal tradeoff ecologically differentiates recently speciated marine bacterioplankton populations. Proc Natl Acad Sci U S A 111(15).

22. Nadell CD, et al. (2013) Cutting through the complexity of cell collectives. Proc R Soc B 280(1755):20122770.

23. Reidl J, Klose KE (2002) *Vibrio cholerae* and cholera: out of the water and into the host. FEMS Microbiol Rev 26(2):125–139.

24. Faruque S, et al. (2006) Transmissibility of cholera: In vivo-formed biofilms and their relationship to infectivity and persistence in the environment. PNAS 103(16):6350–6355.

25. Hayes CA, Dalia TN, Dalia AB (2017) Systematic genetic dissection of chitin degradation and uptake in Vibrio cholerae. Environ Microbiol 19(10):4154–4163.

26. Meibom KL, et al. (2004) The Vibrio cholerae chitin utilization program. Proc Natl Acad Sci U S A 101(8):2524–2529.

27. Drescher K, Nadell CD, Stone HA, Wingreen NS, Bassler BL (2014) Solutions to the Public Goods Dilemma in Bacterial Biofilms. Curr Biol 24(1):50–55.

28. Stocker R, Seymour JR (2012) Ecology and physics of bacterial chemotaxis in the ocean. Microbiol Mol Biol Rev 76(4):792–812.

29. Meibom KL, Blokesch M, Dolganov NA, Wu CY, Schoolnik GK (2005) Chitin induces natural competence in Vibrio cholerae. Science (80-) 310(5755):1824–1827.

30. Bari SMN, et al. (2013) Quorum-sensing autoinducers resuscitate dormant Vibrio cholerae in environmental water samples. Proc Natl Acad Sci U S A 110(24):9926–9931.

31. Borgeaud S, Metzger LC, Scrignari T, Blokesch M (2015) The type VI secretion system of Vibrio cholerae fosters horizontal gene transfer. Science (80-) 347(6217):63–67.

32. Blokesch M, Schoolnik GK (2007) Serogroup conversion of Vibrio cholerae in aquatic reservoirs. PLoS Pathog 3(6):733–742.

33. Zhu J, Mekalanos JJ (2003) Quorum sensing-dependent biofilms enhance colonization in Vibrio cholerae. Dev Cell 5(4):647–656.

34. Takemura AF, Chien DM, Polz MF (2014) Associations and dynamics of Vibrionaceae in the environment, from the genus to the population level. Front Microbiol 5.

35. Hu D, et al. (2016) Origins of the current seventh cholera pandemic. Proc Natl Acad Sci U S A 113(48):E7730–E7739.

36. Faruque SM, et al. (2003) Emergence and evolution of Vibrio cholerae O139. Proc Natl Acad Sci 100(3).

37. Fong JCN, Yildiz FH (2007) The rbmBCDEF Gene Cluster Modulates Development of Rugose Colony Morphology and Biofilm Formation in Vibrio cholerae. J Bacteriol 189(6):2319–2330.

38. Beyhan S, Yildiz FH (2007) Smooth to rugose phase variation in Vibrio cholerae can be mediated by a single nucleotide change that targets c-di-GMP signalling pathway. Mol Microbiol 63(4):995–1007.

39. Fong JCN, Syed KA, Klose KE, Yildiz FH (2010) Role of Vibrio polysaccharide (vps) genes in VPS production, biofilm formation and Vibrio cholerae pathogenesis. Microbiology 156(9):2757–2769.

40. Berk V, et al. (2012) Molecular architecture and assembly principles of Vibrio cholerae biofilms. Science (80-) 337(6091):236–239.

41. Teschler JK, et al. (2015) Living in the matrix: assembly and control of Vibrio cholerae biofilms. Nat Rev Micro 13(5):255–268.

42. Drescher K, et al. (2016) Architectural transitions in Vibrio cholerae biofilms at single-cell resolution. Proc Natl Acad Sci:201601702.

43. Nadell CD, Drescher K, Wingreen NS, Bassler BL (2015) Extracellular matrix structure governs invasion resistance in bacterial biofilms. ISME J 9:1700–1709.

44. Fong JC, et al. (2017) Structural dynamics of RbmA governs plasticity of Vibrio cholerae biofilms. Elife 6. doi:10.7554/eLife.26163.

45. Yan J, Sharo AG, Stone HA, Wingreen NS, Bassler BL (2016) Vibrio cholerae biofilm growth program and architecture revealed by single-cell live imaging. Proc Natl Acad Sci U S A 113(36):E5337–43.

46. Absalon C, Van Dellen K, Watnick PI (2011) A Communal Bacterial Adhesin Anchors Biofilm and Bystander Cells to Surfaces. PLoS Pathog 7(8):e1002210.

47. Kierek K, Watnick PI (2003) The Vibrio cholerae O139 O-antigen polysaccharide is essential for Ca2+-dependent biofilm development in sea water. Proc Natl Acad Sci U S A 100(24):14357–14362.

48. Shcherbo D, et al. (2007) Bright far-red fluorescent protein for whole-body imaging. Nat Methods 4(9):741–746.

49. Kikuchi A, et al. (2008) Structural Characterization of a Thiazoline-Containing Chromophore in an Orange Fluorescent Protein, Monomeric Kusabira Orange ^† ‡^. Biochemistry 47(44):11573–11580.

50. Mizunoe Y, Wai SN, Takade A, Yoshida S-I (1999) Isolation and Characterization of Rugose Form of Vibrio cholerae O139 Strain MO10. Infect Immun 67(2).

51. Burd AB, Jackson GA (2009) Particle aggregation. Ann Rev Mar Sci 1:65–90.

52. Bartlett TM, et al. (2017) A Periplasmic Polymer Curves Vibrio cholerae and Promotes Pathogenesis. Cell 168(1–2):172–185.e15.

53. Giglio KM, Fong JC, Yildiz FH, Sondermann H (2013) Structural Basis for Biofilm Formation via the Vibrio cholerae Matrix Protein RbmA. J Bacteriol 195(14):3277–3286.

54. Schluter J, Nadell CD, Bassler BL, Foster KR (2015) Adhesion as a weapon in microbial competition. ISME J 9(1):139–149.

55. Persat A, et al. (2015) The Mechanical World of Bacteria. Cell 161(5):988–997.

56. Yang DC, Blair KM, Salama NR (2016) Staying in Shape: the Impact of Cell Shape on Bacterial Survival in Diverse Environments. Microbiol Mol Biol Rev 80(1):187–203.

57. Young KD (2006) The selective value of bacterial shape. Microbiol Mol Biol Rev 70(3):660–703.

58. Persat A, Stone HA, Gitai Z (2014) The curved shape of Caulobacter crescentus enhances surface colonization in flow. Nat Commun 5. doi:10.1038/ncomms4824.

59. Rossy T, Nadell CD, Persat A (2018) Cellular advective-diffusion drives the emergence of bacterial surface colonization patterns and heterogeneity. bioRxiv:434167.

60. Martínez-García R, Nadell CD, Hartmann R, Drescher K, Bonachela JA (2018) Cell adhesion and fluid flow jointly initiate genotype spatial distribution in biofilms. PLOS Comput Biol 14(4):e1006094.

61. Smith WPJ, et al. (2017) Cell morphology drives spatial patterning in microbial communities. Proc Natl Acad Sci U S A 114(3):E280–E286.

62. Möller J, Luehmann T, Hall H, Vogel V (2012) The Race to the Pole: How High-Aspect Ratio Shape and Heterogeneous Environments Limit Phagocytosis of Filamentous Escherichia coli Bacteria by Macrophages. Nano Lett 12(6):2901–2905.

63. Horvath DJ, et al. (2011) Morphological plasticity promotes resistance to phagocyte killing of uropathogenic Escherichia coli. Microbes Infect 13(5):426–37.

64. Deng Y, Sun M, Shaevitz JW (2011) Direct Measurement of Cell Wall Stress Stiffening and Turgor Pressure in Live Bacterial Cells. Phys Rev Lett 107(15):158101.

65. de Boer PA, Crossley RE, Rothfield LI, Nelson DR, Jun S (1990) Central role for the Escherichia coli minC gene product in two different cell division-inhibition systems. Proc Natl Acad Sci U S A 87(3):1129–33.

66. Smith DE, Babcock HP, Chu S (1999) Single-polymer dynamics in steady shear flow. Science 283(5408):1724–7.

67. Perkins TT, Smith DE, Chu S (1997) Single polymer dynamics in an elongational flow. Science 276(5321):2016–21.

68. Kawale D, et al. (2017) Polymer conformation during flow in porous media. Soft Matter 13(46):8745–8755.

69. Porter RS, Johnson JF (1966) THE ENTANGLEMENT CONCEPT IN POLYMER SYSTEMS Available at: https://pubs.acs.org/sharingguidelines [accessed october 31, 2018].

70. Colwell RR, et al. (2003) Reduction of cholera in Bangladeshi villages by simple filtration. Proc Natl Acad Sci U S A 100(3):1051–5.

71. Colwell RR (1996) Global climate and infectious disease: the cholera paradigm. Science 274(5295):2025–31.

72. Skorupski K, Taylor RK (1996) Positive selection vectors for allelic exchange. Gene 169(1):47–52.

73. Sia SK, Whitesides GM (2003) Microfluidic devices fabricated in poly(dimethylsiloxane) for biological studies. Electrophoresis 24:3563–3576.

74. Weibel DB, DiLuzio WR, Whitesides GM (2007) Microfabrication meets microbiology. Nat Rev Microbiol 5(3):209–218.

75. De Lorenzo V, Timmis KN (1993) Analysis and construction of stable phenotypes in gram-negative bacteria with Tn5-and Tn10-derived minitransposons. Methods Enzymol 235:386–405.

76. Tacket CO, et al. (1995) Initial Clinical Studies of CVD 112 Vibrio cholerae O139 Live Oral Vaccine: Safety and Efficacy against Experimental Challenge. J Infect Dis 172(3):883–886.

